# Elucidating the *Staphylococcus aureus* TSST-1 regulatory network as a response to vaginal pH

**DOI:** 10.1101/2025.09.06.674626

**Authors:** Carla S. Maduta, Nicholas R. Walton, Emily P. Barbosa, Erica N. DeJong, Karine Dufresne, John K. McCormick

## Abstract

Menstrual toxic shock syndrome (mTSS) is a life-threatening disease caused by the *Staphylococcus aureus* superantigen TSST-1. At menstruation, the typical acidic vaginal environment rises to near neutral pH, which allows for optimal TSST-1 production. However, the regulation network which alters toxin production in response to pH is largely unknown, despite the importance of this cue in the vaginal environment. To mimic the vaginal environment, we used Vaginally Defined Medium to assess TSST-1 promoter (*tst*) activity in the mTSS strain *S. aureus* MN8 and discovered a significant upregulation of *tst* expression occurring at pH 4.5 in low glucose environment, referred to as the ‘acidic virulence surge’. This increase was also observed in all the regulatory mutant backgrounds tested, including in the absence of *saeS*, which has previously been thought to be required for TSST-1 production. Recent studies in non-mTSS *S. aureus* strains have identified GraXRS as a pH sensor, in addition to its function in cationic antimicrobial peptide sensing. We therefore hypothesized that GraXRS alters TSST-1 expression at low pH. Deletion of the sensor-kinase *graS* resulted in the loss of TSST-1 surge at pH 4.5, indicating that GraXRS is required for the acidic virulence surge. We also found expression from the SaeRS P1 and SarA promoters to be significantly attenuated in the Δ*graS* background. At low pH, the absence of GraS resulted in the least amount of T cell activation from *S. aureus* supernatants when compared to other regulatory mutants, suggesting that GraXRS is the dominant activator at pH 4.5. Finally, we developed an *in vivo* murine model to measure *tst* expression using luciferase expression. Our results demonstrate a complex sequence of events that occur in response to changes in pH and further suggests that GraXRS is the main activator of TSST-1 at low pH in *S. aureus*.

**AUTHOR SUMMARY:** Menstrual toxic shock syndrome is a life-threatening hyperinflammatory disease, resulting from the production of a toxin named TSST-1 by *Staphylococcus aureus*. Environmental cues within the vagina are sensed by *S. aureus*, resulting in changes in the production of TSST-1. A key environmental cue present within the vagina is acidic pH, which has previously been thought to limit TSST-1 production. Here, we use a luminescent reporter assay to determine how pH affects expression of TSST-1 in a vaginal mimicking medium. We found that expression of the toxin drastically increases at a pH of 4.5, and key TSST-1 regulators are not responsible for this phenotype. We found that deletion of *graS* abolishes the toxin’s production at pH 4.5 and limits the activation of primary T cells. We also established an *in vivo* model of murine vaginal colonization to examine the expression of TSST-1. Our results highlight the ability of TSST-1 to be produced outside of conventional menstrual conditions and provides insight on the necessity of modeling the environment when studying bacterial virulence factors.

## INTRODUCTION

*Staphylococcus aureus* is a highly prolific human colonizer and opportunistic pathogen. *S. aureus* is often found on mucosal surfaces including the vagina, where it can be part of the commensal microbiota, or take part in dysbiosis and pathogenesis [1]. The most highly researched pathology attributed to vaginal *S. aureus* is the life-threatening disease known as menstrual toxic shock syndrome (mTSS). The development of mTSS is caused by the production of the superantigen toxic shock syndrome toxin-1 (TSST-1). In essence, TSST-1, like other superantigens, is able to bypass the peptide specificity required in T cell activation [2]. Superantigens crosslink MHC-II molecules on antigen presenting cells with T cell receptors, leading to the activation of numerous T cells. This can initiate a pro-inflammatory cytokine storm that may lead to toxic shock syndrome that can be fatal if left untreated.

The production of TSST-1 is tightly controlled by an array of staphylococcal regulators which respond to environmental cues, many of which are two-component systems (TCS) (**Fig. 1**). *S. aureus* encodes 16 TCS, each composed of a sensor histidine kinase that signals to its response regulator to initiate changes in transcription [3]. Several TCS have been implicated in TSST-1 regulation including Agr, SrrAB and SaeRS. The Agr system responds to cell density and functions as an indirect activator of TSST-1 by relieving repression by Rot [4]. The SrrAB system is an activator of TSST-1 when oxygen levels are increased, such as through the insertion of menstrual devices [5]. Finally, SaeRS is believed to be required for TSST-1 production, as its deletion or inhibition eliminate TSST-1 production [6,7]. While SaeRS is the dominant TSST-1 activator, it is unclear what the sensor is detecting, although it is hypothesized to detect membrane perturbations [8]. Other stand-alone regulators have been implicated in TSST-1 regulation, with the most notable being CcpA which represses TSST-1 transcription in response to high glucose levels that are persistent outside of menstruation [9]. SarA has been previously identified as a repressor in the *S. aureus* strain MN8 [6], yet in Vaginally Defined Medium (VDM), it can function as an activator [7]. Members of the vaginal microbiota can also alter the expression of TSST-1 *in vitro* by manipulating *S. aureus* regulation systems [10].

**Figure 1.**
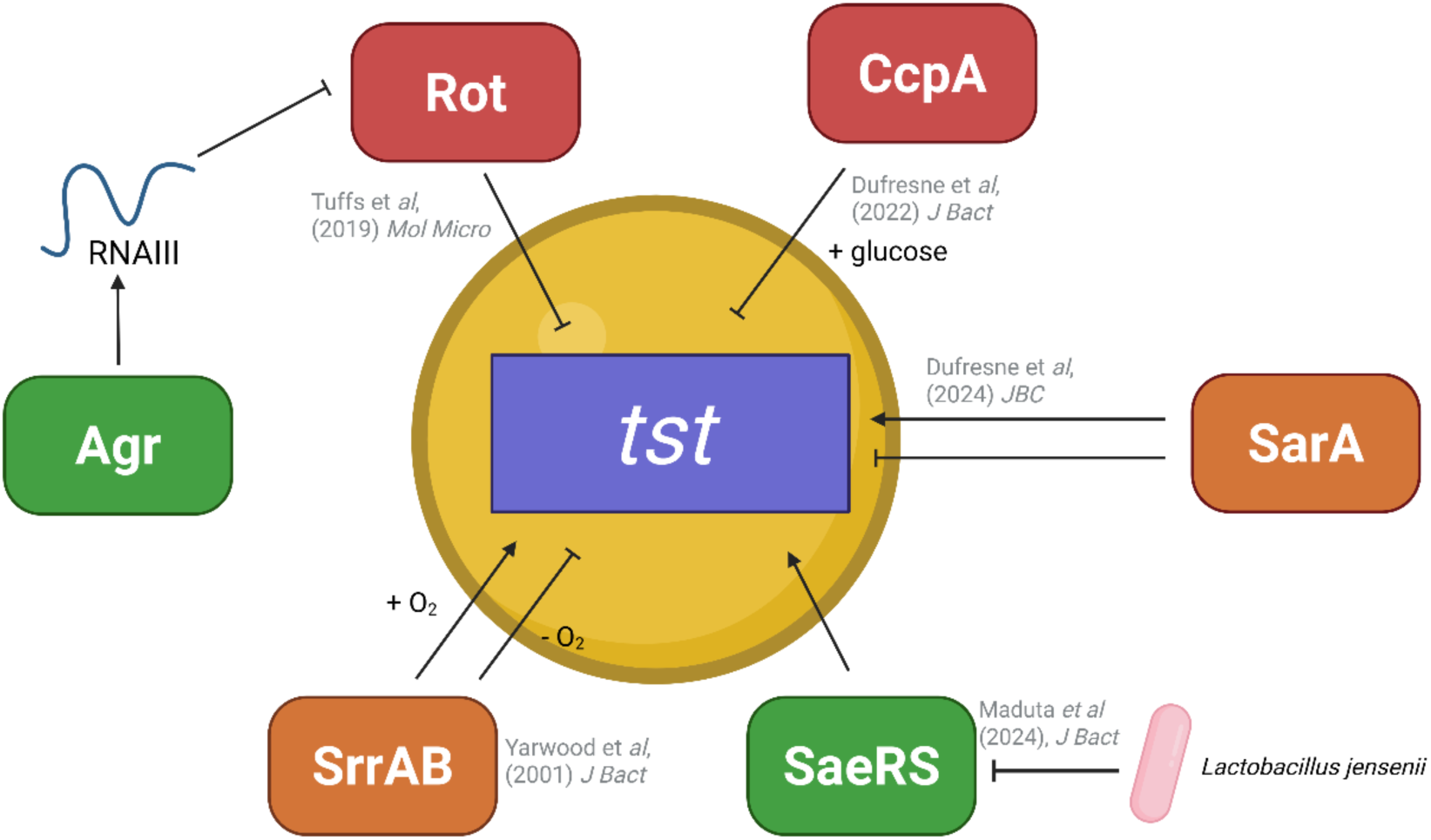
Regulation of *tst* expression in *S. aureus* MN8. Activators of *tst* expression are shown in green, repressors in red, and those with mixed regulatory activity in orange. SaeRS is the direct and dominant activator of TSST-1 production. SarA functions as an activator in Vaginally Defined Medium in strain MN8, although activity differs in other environments and genetic backgrounds. SrrAB activates TSST-1 production in response to oxygen. Agr indirectly activates *tst* expression by relieving Rot repression. CcpA represses in response to glucose. Figure made with Biorender.com.

A critical defense system within the vagina is acidic pH, via the production of lactic acid by a variety of *Lactobacillus* species [11–13]. Typically, vaginal pH ranges from 3.5–4.5 [14,15], which serves as a protective function by limiting the growth of vaginal pathogens, which may include *S. aureus*. It is has long been postulated that acidic pH represses TSST-1 expression as initial studies were unable to detect toxin production below a pH of 6.5 [16–19]. This pH effect contributes to the restriction of mTSS to menstruation, which is when vaginal pH rises from acidic to near neutral levels. Despite the importance of pH as an environmental cue in the vagina, the *S. aureus* signaling pathway which responds to pH and regulates TSST-1 expression has not been identified. Recently, in non-mTSS *S. aureus* strains, the TCS GraXRS was identified to have a role in bacterial survival in acidic environments [20–22]. Here, we set out to elucidate the TSST-1 regulatory network as a response to pH changes. Using phenotypic assays to determine the expression of TSST-1, we identified GraXRS to be the predominant positive regulator of the toxin’s expression at acidic pH.

## RESULTS

### An abrupt increase in TSST-1 expression and production occurs at pH 4.5

While pH has been considered a critical environmental cue in TSST-1 expression, most transcriptional studies have been performed in standard laboratory media which may not necessarily represent the vaginal environment. To best mimic mTSS conditions, we used VDM adjusted to 0.7 mM glucose in order to relieve CcpA mediated repression [9,10,23,24]. VDM was adjusted to pHs from 3.0–8.0 and a luciferase reporter assay used as a measure of activity from the *tst* promoter (**Fig 2A-J**). In alignment with previous observations [19], optimal *tst* expression occurred at pH 6.8 (**Fig. 2H**), which is a pH level consistent with menstruation. As pH decreased, *tst* expression decreased drastically (**Fig. 2K**) although surprisingly, at pH 4.5 there was a large increase in *tst* expression (**Fig. 2D, 2K**), herein referred to as the “acidic virulence surge.” This pH level is the most acidic at which *S. aureus* was able to grow (**Fig. 2A-D**). The transcriptional data was further confirmed via anti-TSST-1 Western blotting where TSST-1 production stopped being detected at pH 5.0 and returned at pH 4.5 (**Fig. 2L**). Overall, these data confirm that acidic pH is generally repressive in a vaginal-like environment yet simultaneously identify a potentially alarming increase in virulence at a growth-limiting pH.

**Figure 2.**
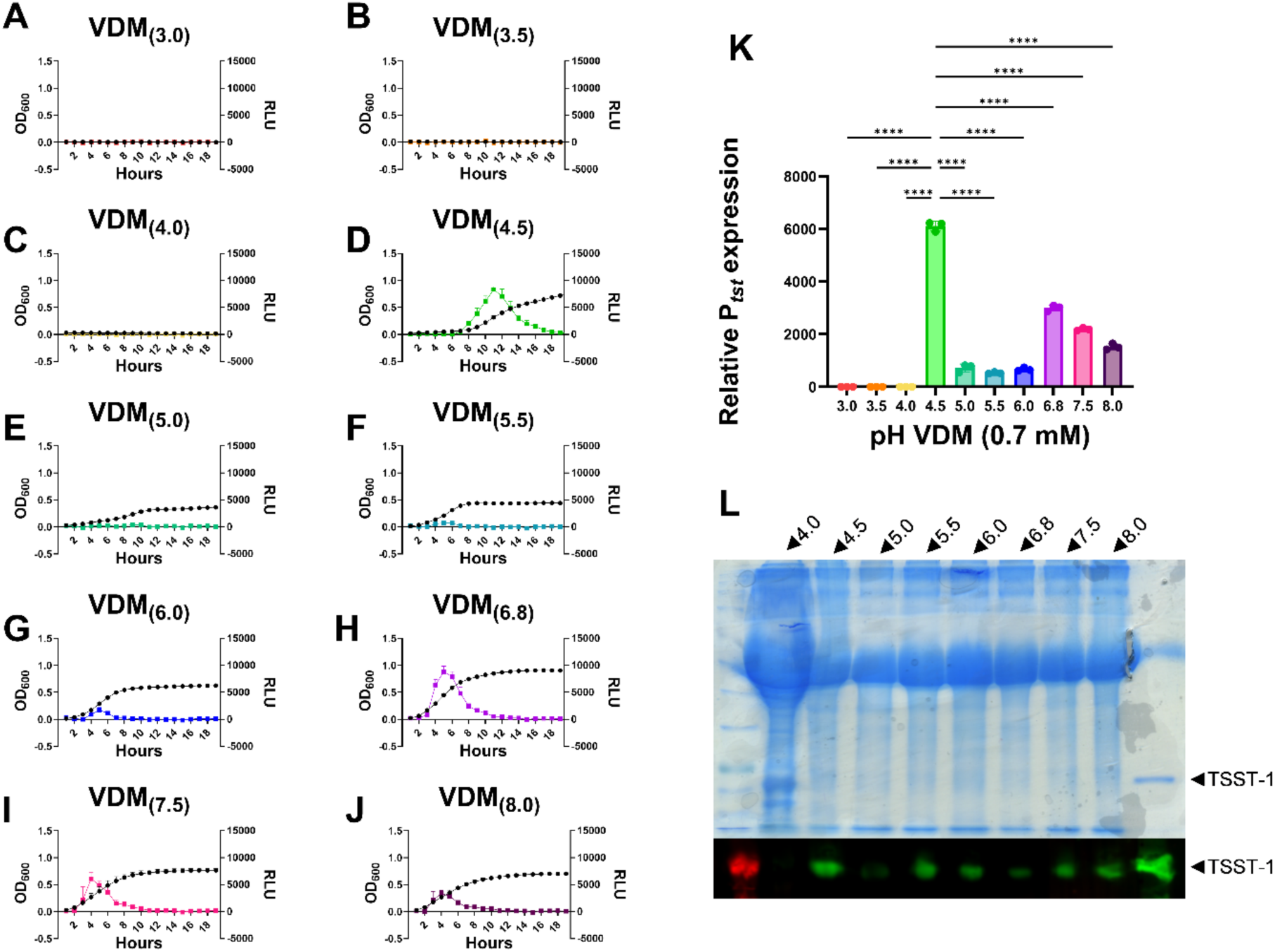
Expression and production of TSST-1 increases significantly at pH 4.5. Wild-type MN8 containing pAmilux::p*tst* was grown in VDM (0.7 mM glucose) from pHs 3.0-8.0 (A-J) and readings were taken every hour. Relative expression (K) was determined as the area under the curve (AUC) of luminescence over the AUC of growth. Wild-type MN8 was also grown overnight, standardized to 12 OD units, followed by SDS-PAGE and anti-TSST-1 Western blotting (L). Statistical analysis was performed with GraphPad Prism 10 using a One-Way ANOVA (* P≤0.05, ** P≤0.01, *** P≤0.001, **** P≤0.0001).

### Acidic activation does not overcome glucose repression

While pH is an environmental cue which fluctuates greatly at menstruation, glucose levels also decrease significantly at menstruation. We have previously demonstrated that TSST-1 production does not occur in the presence of high glucose due to CcpA repression [9]. However, it is unknown whether the surge in TSST-1 production at pH 4.5 can be overridden by glucose repression. Therefore, we set out to investigate whether pH or glucose is the dominant regulatory cue at pH 4.5. A luciferase assay was performed in high glucose VDM (60 mM) and *tst* expression from wild-type MN8 did not occur at any pH level, nor was TSST-1 detected by Western blotting (**Fig. 3**). These data indicate that between acidic driven activation or glucose driven repression, glucose is the dominant cue in the vaginal environment.

**Figure 3.**
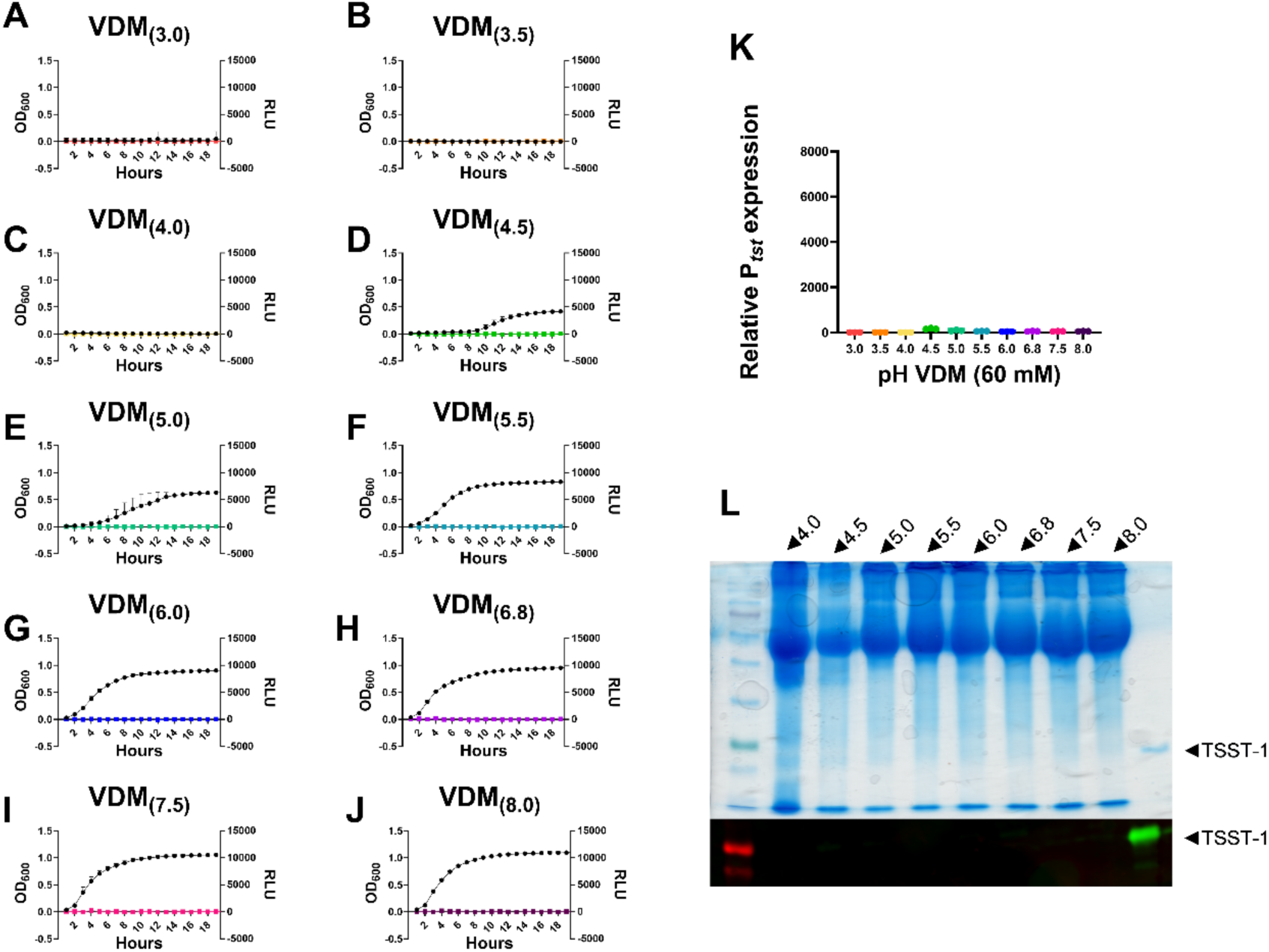
Glucose repression of *tst* overrides acidic pH activation. Wild-type *S. aureus* MN8 containing pAmilux::p*tst* was grown in VDM containing 60 mM glucose from pHs 3.0-8.0 and readings were taken every hour. Relative expression (K) was determined as the area under the curve (AUC) of luminescence over the AUC of growth. Wild-type *S. aureus* MN8 was also grown overnight, standardized to 12 OD units, followed by SDS-PAGE and anti-TSST-1 Western blotting (L). Statistical analysis was performed with GraphPad Prism 10 using a One-Way ANOVA.

### Known TSST-1 regulators are not responsible for the acidic virulence surge

To investigate the surge in TSST-1 production occurring in low glucose conditions (**Fig. 2D, 2K**), we turned to a plethora of regulatory mutants which were previously only tested at pH 6.8 [7]. If any of these regulatory systems were responsible for the increase, we expected to observe no increase in *tst* expression at pH 4.5. The luciferase assay was repeated in previously constructed *tst* luminescence reporters in the following mutational backgrounds: Δ*ccpA,* Δ*saeS,* Δ*sarA,* Δ*srrAB*, Δ*agr,* Δ*rot*, and Δ*agr*Δ*rot*.

In the Δ*agr* background, no *tst* promoter activity was observed given that repression from Rot is maintained (**Fig. 4A**) [4]; yet, TSST-1 was still detectable via Western blotting (**Fig. 4B, 4C**). In the Δ*rot* and Δ*agr*Δ*rot* backgrounds, TSST-1 was expressed to similar levels as the wild-type. When examining the difference between expression at pH 6.8 and pH 4.5 in these regulatory mutants, the strains phenocopied the wild-type and demonstrated the acidic virulence surge. Therefore, these data suggest the Agr-Rot axis is not responsible for the low pH phenotype as elimination of this axis maintained the response.

**Figure 4.**
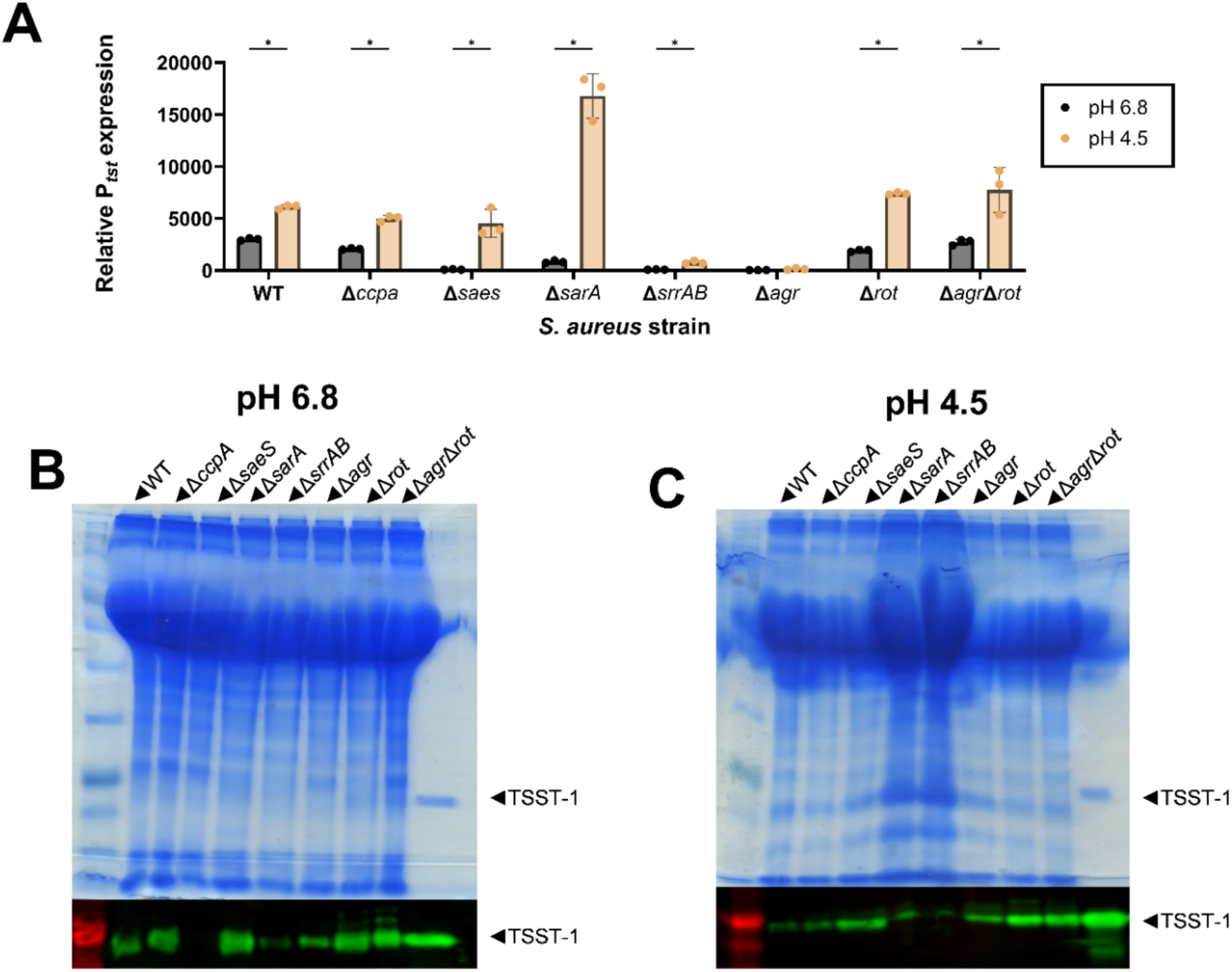
Regulatory mutants display a surge in TSST-1 expression at pH 4.5. Regulatory deletion mutants containing pAmilux::p*tst* were grown in VDM at pHs 6.8 and 4.5 (A) and readings were taken every hour. Relative expression was determined as the area under the curve (AUC) of luminescence over the AUC of growth. The wild-type luminescence data was used as a comparison reference and is the same as in Fig. 2K. The mutants were also grown overnight, standardized to 12 OD units, followed by SDS-PAGE and anti-TSST-1 Western blotting; pH 6.8 (B) and pH 4.5 (C). Statistical analysis was performed with GraphPad Prism 10 using multiple unpaired t tests (* P≤0.05).

Given that SaeRS is the major activator of TSST-1 expression, we next sought to investigate whether this regulatory system was responsible for the spike at pH 4.5. In the Δ*saeS* background we observed no *tst* expression (**Fig. 4A**) nor production (**Fig. 4B**) at pH 6.8, which is consistent with SaeRS being required for TSST-1 expression [6]. However, at pH 4.5, activity from the promoter was surprisingly detected, as was the TSST-1 protein via Western blotting (**Fig. 4A, 4C**) which suggests that SaeRS is not responsible for this spike.

We next turned to the *S. aureus* MN8 Δ*srrAB* background, as SrrAB is another activator of TSST-1 production [5]. Nevertheless, the surge at pH 4.5 was modestly detected, ruling out this regulatory system (**Fig. 4A**). Similarly, the Δ*ccpA* strain also displayed increased expression at pH 4.5. Finally, we turned to Δ*sarA*, as the regulator was recently established to be a TSST-1 activator in VDM [7]. The Δ*sarA* strain also demonstrated increased *tst* expression at pH 4.5, ∼3-fold larger than the wild-type (**Fig. 4A**). Altogether, these results indicate that previously identified TSST-1 regulators are not responsible for the increase in *tst* expression at pH 4.5 and there must be an additional positive regulator which acts in acidic conditions.

### GraXRS expression is enhanced at acidic pH

Previous studies from Flannagan et al. [20] and Kuiack et al. [21] identified a novel function for the TCS GraXRS in survival and genetic regulation at acidic pH. Tn-seq experiments using the Nebraska Transposon Library also identified *graS* as a key gene required for the bacterial stress response in TSB at pH 4.5 vs 7.3 [25]. We therefore hypothesized that GraXRS may play a role in regulating TSST-1 production in response to pH.

Given that expression of regulatory systems differs among *S. aureus* strains, as well as across different media, we first aimed to determine whether GraXRS was adequately expressed in *S. aureus* MN8 when grown in VDM. We constructed a luciferase reporter for the GraXRS promoter and found that activity was highest at acidic pH and significantly decreased towards neutral and basic pH ranges (**Fig. 5K**). These results indicate that GraXRS expression is altered as a response to acidic pH in *S. aureus* MN8.

**Figure 5.**
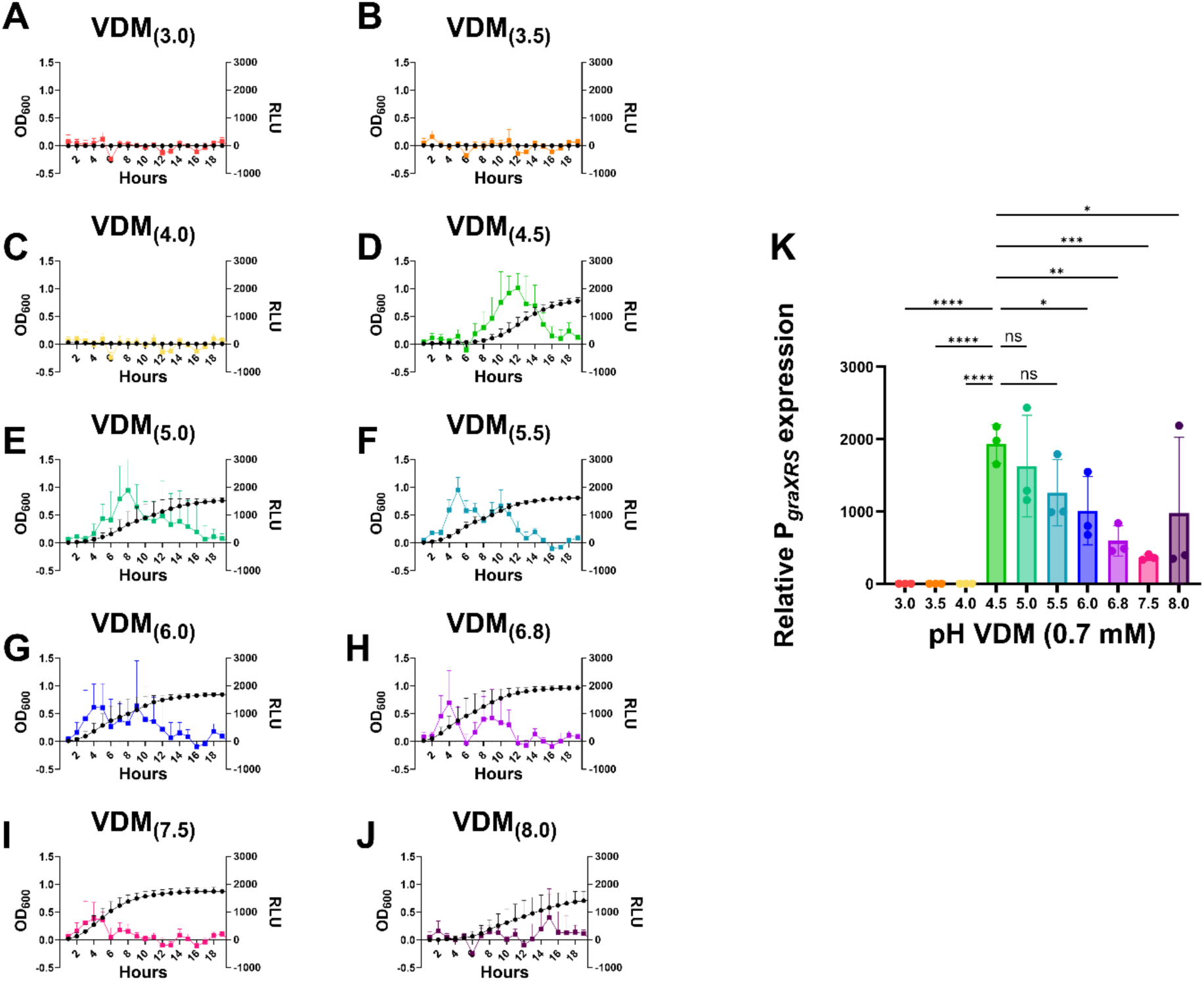
Expression from the *graXRS* promoter is enhanced at acidic pH. Wild-type MN8 containing pAmilux::p*graXRS* was grown in low glucose VDM from pHs 3.0-8.0 (A-J) and readings were taken every hour. Relative expression (K) was determined as the area under the curve (AUC) of luminescence over the AUC of growth. Statistical analysis was performed with GraphPad Prism 10 using a One-Way ANOVA (* P≤0.05, ** P≤0.01, *** P≤0.001, **** P≤0.0001).

### GraXRS is required for TSST-1 production at pH 4.5

As we established that GraXRS was expressed by *S. aureus* MN8 in VDM, we next generated a clean deletion of the sensor kinase *graS* gene to investigate how the deprivation of this TCS altered TSST-1 expression. A clean deletion of *graS* was constructed, followed by electroporation with pAmilux::p*tst*, as well as the reporter pAmilux plasmids for *sae* (P1)*, sarA*, and *agr* (P3). When testing the Δ*graS* strain for the production of TSST-1, the toxin could be detected by Western blot between pH ranges 5.0 to 8.0 (**Fig. 6A**). Importantly, at a pH of 4.5, TSST-1 was not detected. Relative to the wild-type *S. aureus* MN8, Δ*graS* expressed significantly less *tst* between pH ranges 4.5 to 8.0 (**Fig. 6B**), suggesting that GraXRS is a positive regulator of TSST-1. To further validate the role of GraXRS, a luciferase assay was performed with wild-type *S. aureus* MN8 in the presence of the GraR inhibitor MAC-545496 at a pH of 4.5 [26]. While did note some growth inhibition with as little as 0.25 µg/ml of MAC-545496, inhibitory effects on *tst* expression were observed beyond 0.5 µg/ml (**Fig. S1**). Despite the effects on growth, inhibition of GraR provides further evidence into the role of this two-component system at pH 4.5.

**Figure 6.**
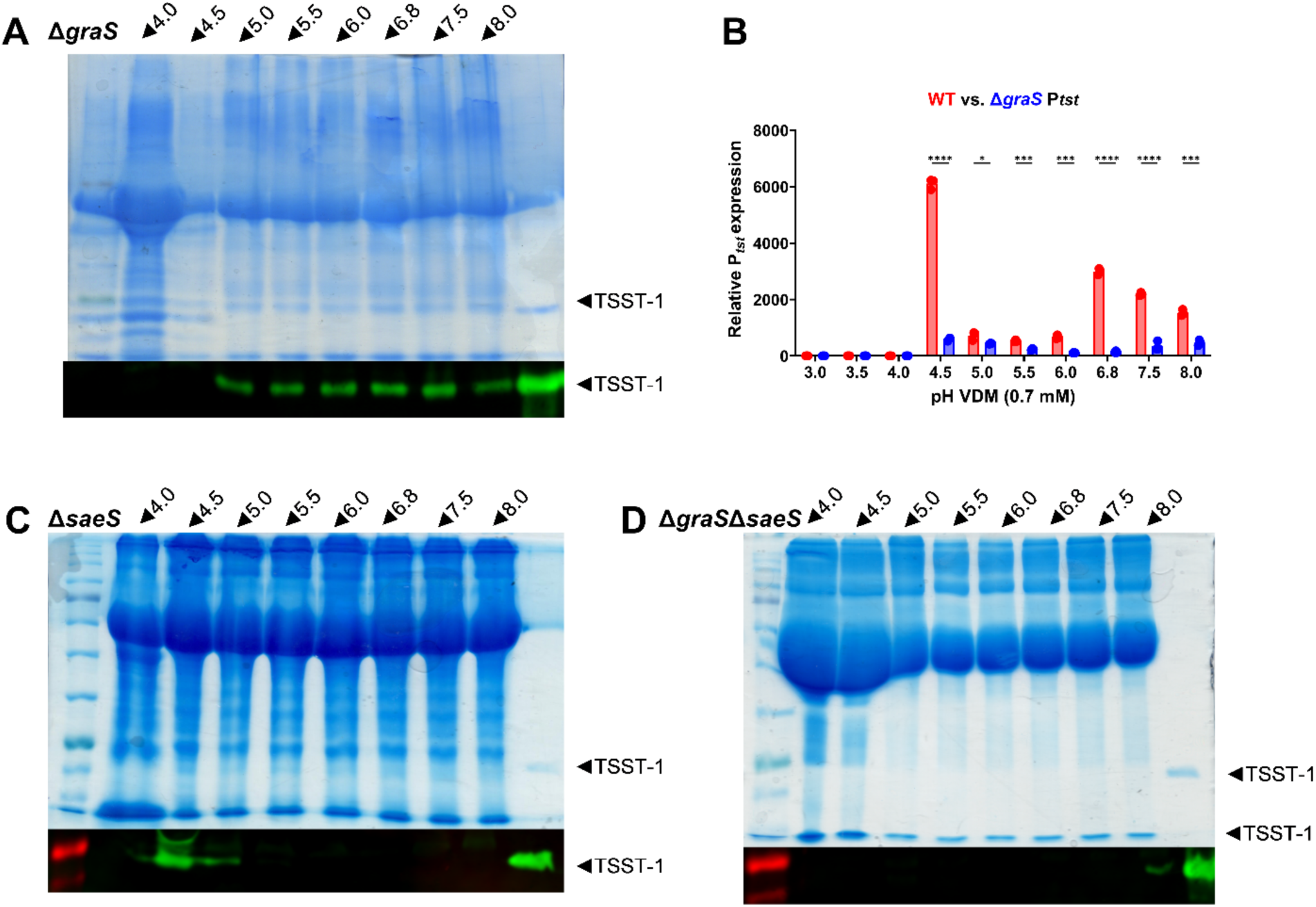
GraXRS is an activator of TSST-1 expression at pH 4.5. MN8 Δ*graS* (A), Δ*saeS* (C) and Δ*graS*Δ*saeS* (D) were grown in VDM from pHs 4.0-8.0 overnight, standardized to 12 OD units, followed by SDS-PAGE and anti-TSST-1 Western blotting. Wild-type MN8 or MN8 Δ*graS* containing pAmilux::p*tst* were grown in VDM from pHs 3.0-8.0 (B) and readings were taken every hour. Relative expression was determined as the area under the curve (AUC) of luminescence over the AUC of growth. The wild-type luminescence data is used as a comparison reference and is the same as in Fig. 2K. Statistical analysis was performed with GraphPad Prism 10 using a One-Way ANOVA (* P≤0.05, ** P≤0.01, *** P≤0.001, **** P≤0.0001).

Considering that the acidic virulence surge was detected both in wild-type S. aureus MN8 as well as in the absence of various regulatory systems (**Fig. 4**), we sought to investigate whether the deletion of *graS* would also reverse the increase in a non-WT background. Given the surprising production of TSST-1 by MN8 Δ*saeS* (**Fig. 4D**, **Fig. 6C**), we created a double Δ*graS* Δ*saeS* mutant. In this strain lacking both *graS* and *saeS,* TSST-1 was no longer detected at a pH 4.5 (**Fig. 6D**). A faint band was detected at pH 8.0, however this is not a biologically relevant vaginal pH. Taken together, the reversal of the acidic virulence surge indicates that GraXRS is required for activation of TSST-1 production at pH 4.5.

Given that *tst* expression was affected at other pH levels (**Fig. 6B**), yet *graXRS* promoter activity was decreased (**Fig. 5**), this suggests that GraXRS may not be responsible for the decreased expression at pH ranges above 4.5. This led us to hypothesize that other positive regulatory systems may be implicated. Examining the P3 promoter from *agr*, we saw similar expression in wild-type *S. aureus* MN8 and MN8 Δ*graS* backgrounds, suggesting that differential regulation of *agr* is not responsible for the changes in *tst* expression (**Fig. 7A**). While there appears to be a large increase in relative P3 expression at pH 4.5 (**Fig. 7A**), this is likely due to a 2-to-3-hour lag in growth in the Δ*graS* background relative to the wild-type, affecting the area under the OD600 curve, and not due to increased promoter activity (**Fig. S2**). However, when examining activity from the P1 promoter from *sae* (**Fig. 7B**) and the *sarA* promoter (**Fig. 7C**), there were significant decreases in the Δ*graS* background, following the trend in *tst* expression (**Fig. 6B**). Overall, these data indicate that GraXRS affects other regulatory systems, but it is only required for TSST-1 production at pH 4.5.

**Figure 7.**
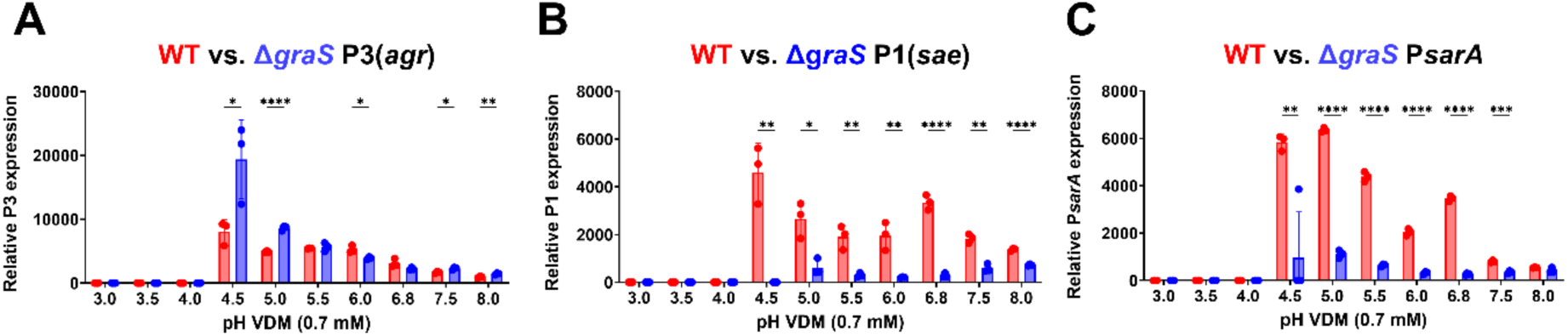
Expression from the *sae* P1 and *sarA* promoters are significantly decreased in MN8 Δ*graS*. Wild-type MN8 or MN8 Δ*graS* containing (A) pAmilux::p3*(agr)*, (B) pAmilux::p1*(sae)*, or (C) pAmilux::p*sarA* were grown in VDM from pHs 3.0-8.0 and readings were taken every hour. Relative expression was determined as the area under the curve (AUC) of luminescence over the AUC of growth. Statistical analysis was performed with GraphPad Prism 10 using a One-Way ANOVA (* P≤0.05, ** P≤0.01, *** P≤0.001, **** P≤0.0001).

### Acidic pH significantly decreases T cell activation by S. aureus

To investigate whether the differences in *tst* expression and TSST-1 production among the regulatory mutant backgrounds corresponded to a biological difference, a T cell activation assay was performed using human peripheral blood mononuclear cells (PBMCs). Briefly, *S. aureus* strains were grown in VDM at either pH 6.8 or pH 4.5, and the supernatants were then exposed to PBMCs from healthy human blood donors for 18h, followed by an interleukin-2 (IL-2) ELISA to assess T cell activation. To make quantitative comparisons of toxin potency among the strains, the 1.6×10^−4^ dilution data points were normalized to the wild-type at pH 6.8 (**Fig. 8D**). A bacterial supernatant eliciting greater IL-2 production at higher dilution levels is considered to have greater toxin potency.

**Figure 8.**
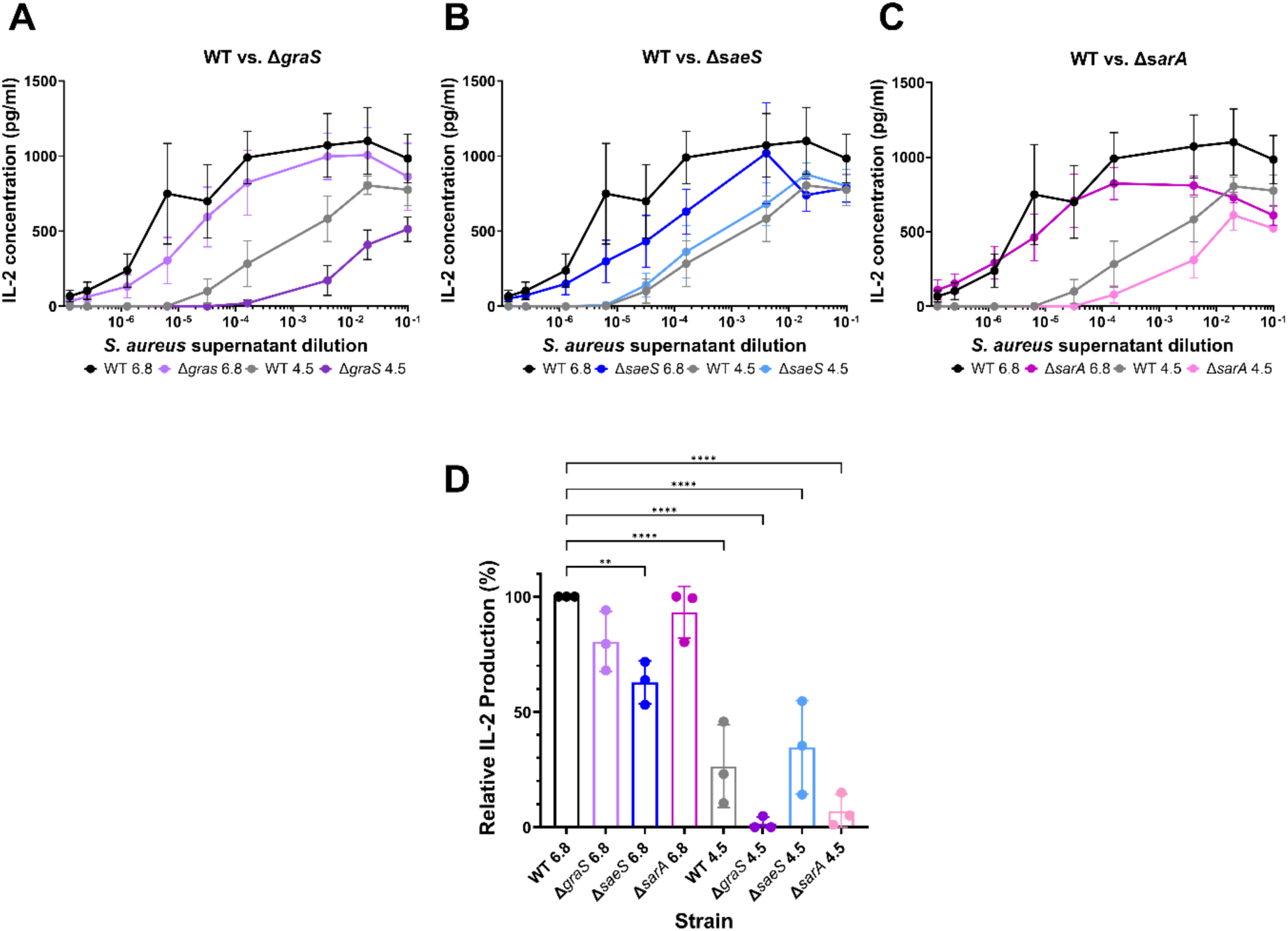
Acidic pH significantly decreases T cell activation by *S. aureus*. Wild-type MN8 and deletion mutants were grown in VDM at pH 6.8 or 4.5 and the supernatants were collected. Human blood was collected from three healthy donors and the PBMCs were exposed to the *S. aureus* supernatants. Cell supernatants were then used in an IL-2 ELISA as a measure of T cell activation. Averages of IL-2 from the three donors were plotted (A-C), and the 1.6×10^−4^ dilution data points were standardized to the wild-type at pH 6.8 to compare potency (D). The wild-type data is used as a comparison reference and is the same across panels A-C. Statistical analysis was performed in GraphPad Prism 10, using a one-way Anova (** P≤0.01, **** P≤0.0001).

At pH 6.8, there was no significant difference in IL-2 production between wild-type *S. aureus* MN8 and Δ*graS* (**Fig. 8A**) or Δ*sarA* (**Fig. 8C**) strains (**Fig. 8D**). When comparing the wild-type to the Δ*saeS* condition (**Fig. 8B**), there was a significant decrease in IL-2 production (**Fig. 8D**), which was expected due to the decreased TSST-1 production from this mutant. Lowering the pH to 4.5 also resulted in a significant decrease in IL-2 production across all strains in comparison to the wild-type at pH 6.8 (**Fig. 8D**). All regulatory mutants had higher IL-2 production at pH 6.8 than at 4.5, with the Δ*graS* condition demonstrating the least amount of T cell activation in low pH. This data suggests greater superantigen activity at pH 6.8 than at pH 4.5 grown *S. aureus*, with the least amount of activation occurring in the absence of GraS. Overall, this data indicates that GraXRS is the dominant activator at pH 4.5.

### GraS and SaeS are activators of tst expression in vivo

To date, a plethora of studies have been done *in vitro* to examine the regulation of TSST-1 production and identify key regulators (**Fig. 1**). However, there is little to no knowledge regarding the relevance of these regulators in an *in vivo* model of *S. aureus* vaginal colonization. pAmilux has been used primarily for *in vitro* studies, as well as an *ex vivo* study on *srr* expression in hypoxic skeletal tissues [27]. Adapting our laboratory’s vaginal colonization workflow [28], we set out to develop a model for examining *tst* promoter activity *in vivo*, and to determine the contributions of *graS* and *saeS* in the neutral pH mouse vagina.

Female BALB/c mice (7-9 weeks old) were inoculated with 1×10^6^ CFU of wild-type *S. aureus* MN8, or the Δ*graS* and Δ*saeS* mutants, each harboring pAmilux::p*tst*, and imaged from 6 to 24 hours post-infection (hpi). Generally, *tst* promoter activity was observed starting from 8 hours (**Fig. 9A, 9B**). The wild-type strain resulted in the most *tst* promoter activity at 12 and 24 hpi among the three strains (**Fig. 9C, 9D**). The Δ*saeS* group produced significantly less light than the wild-type *S. aureus* MN8 group at both timepoints, while the Δ*graS* group only trended downwards at 24 hours hpi (**Fig. 9D**). Taken together, these data support that GraRS (24 hpi) and SaeRS (8-24 hpi) are activators of TSST-1 production *in vivo*, with SaeRS being required for *tst* expression in a neutral pH vaginal colonization model.

**Figure 9.**
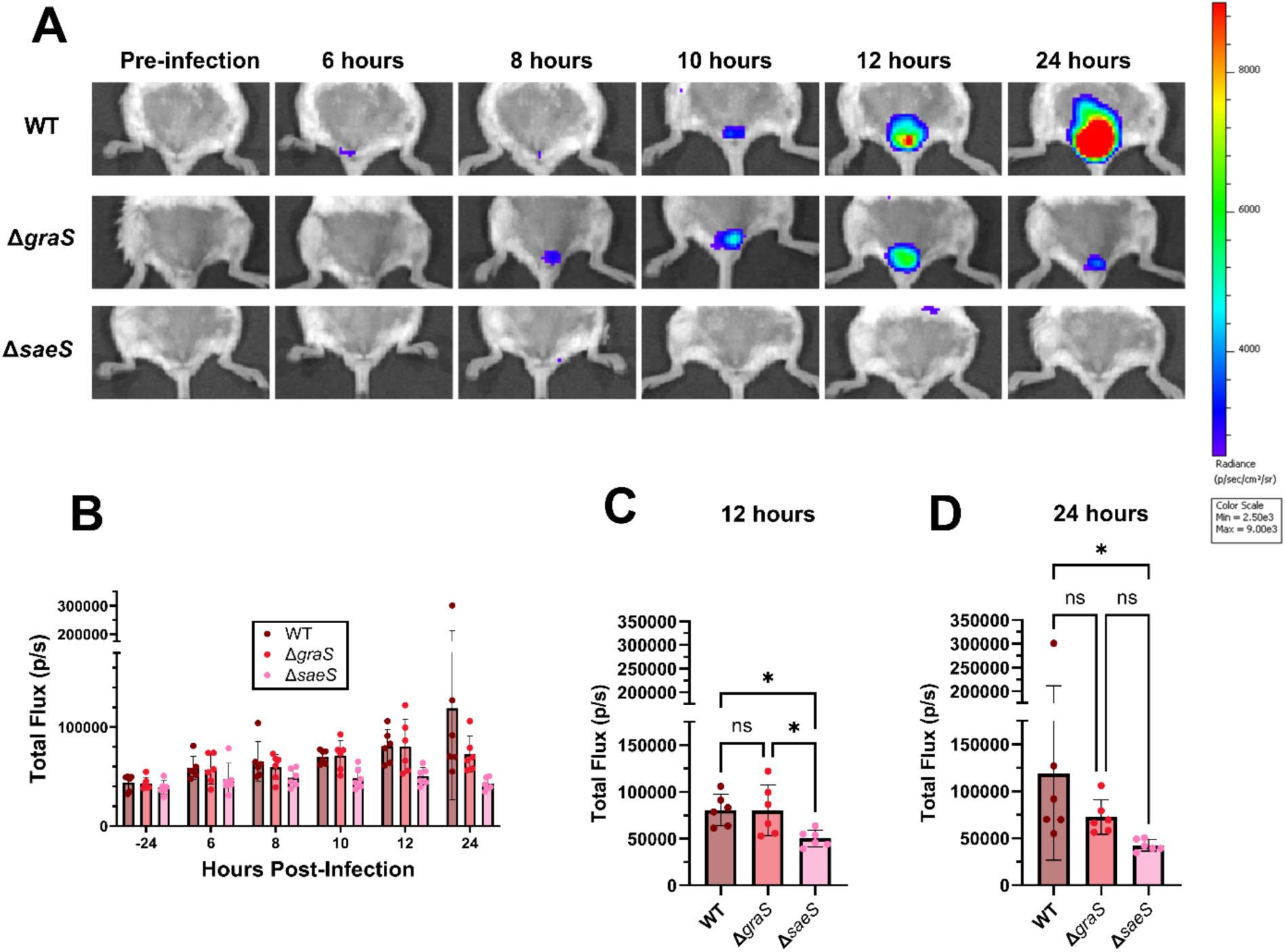
GraRS and SaeRS are activators of *tst* promoter expression in a neutral pH vaginal colonization model. 7–9-week-old BALB/c mice (Jackson Labs) (n=6) were pre-treated with chloramphenicol water *ad libitum* prior to infection with genetically modified *S. aureus* MN8 strains harboring pAmilux::p*tst*. Mice were imaged using *In Vivo* Imaging System (IVIS) at 6, 8, 10, 12 and 24 hpi. One representative mouse is shown for each bacterial group (A). The region of interest (vagina) was identified, and total flux (photons/second) was plotted (B, C, D). Statistical analysis was performed on panels C and D with GraphPad Prism 10 using a One-Way ANOVA (* P≤0.05).

## DISCUSSION

The environmental cues present in the vagina play a critical role in the production of TSST-1 by *S. aureus*. It has long been suggested that the optimal environment in which TSST-1 is produced aligns with the conditions seen at menstruation: decreased glucose, increased oxygen, and near-neutral pH [16]. While the regulation network which alters TSST-1 production has been extensively studied, the role of pH as a regulatory stimulus has remained elusive. In this study, we demonstrate that acidic pH generally reduces TSST-1 production, but at the outer boundary of growth, production of TSST-1 is exaggerated.

Through testing TSST-1 production at pHs ranging from 3.0–8.0, we determined that optimal expression occurs at ∼ pH 6.8 (**Fig. 2**). Interestingly, we characterized a new window of expression at pH 4.5, occurring at the lowest pH at which *S. aureus* MN8 could grow. This increased expression was evident in all the regulatory mutant backgrounds at varying magnitudes (**Fig. 4**), except for in the absence of *graS* (**Fig. 6**), suggesting that the TCS GraXRS activates expression of the toxin at this pH level. An additional key finding from this work is the ability for TSST-1 to be produced in *S. aureus* MN8 Δ*saeS* at pH 4.5 (**Fig. 6C**). Our previous work identified SaeRS as required for the production of TSST-1 [6], which we have now established to be pH-dependent. SaeRS is well known to be vastly responsible for virulence gene expression [29] and without it many key virulence determinants, such as α-toxin [30], are not expressed. Consequently this system is a common target in the development of anti-virulent compounds [7,31]. It is unknown whether this acidic virulence surge we have observed occurs with other SaeRS regulated genes. It would be interesting to investigate other toxins in this context, especially as they relate to other niches known to be exposed to acidic conditions, such as the skin. Given that expression from the Sae P1 promoter was decreased in the absence of GraS (**Fig. 7B**) and the double Δ*graS* Δ*saeS* mutant did not display the acidic virulence surge (**Fig. 6D**), anti-virulence strategies should also further consider targeting the GraXRS system when considering SaeRS regulated genes. While our work suggests there could be overlapping signaling between GraXRS, SaeRS, and SarA (**Fig. 7**), we were unable to detect the GraR binding site upstream of these systems [32]. It is therefore likely that an unknown intermediary is involved.

While the increased virulence we have detected is worrisome, an important finding from this work is that glucose repression overrides pH activation (**Fig. 3**). The vagina is an acidic environment with high glucose and glycogen levels. Estrogen is responsible for the maturation of vaginal epithelial cells and the buildup of glycogen [33], which is then catabolized by amylases into smaller sugars [34–37]. These sugars are then used by lactobacilli, such as *Lactobacillus crispatus*, to form either D- or L-lactic acid to acidify the environment. With a lactobacilli-dominated microbiota, a vaginal pH of 4.5 should be associated with abundant glucose levels. The high glucose levels would therefore prevent the acidic virulence surge from occurring (**Fig. 3**). However, as only an abundance of lactobacilli are associated with higher glycogen levels [12,38], their decreased abundance, such as in vaginal dysbiosis, may alter sugar dynamics. Therefore, vaginal dysbiosis may be a setting in which glucose levels are drastically altered, and repression of TSST-1 can be relieved, allowing for the acidic virulence surge we have observed to occur. This disturbance would be particularly worrisome as *Gardnerella vaginalis* and *Lactobacillus iners* are two bacteria associated with dysbiosis which can also interact with *S. aureus* and exacerbate TSST-1 production [10]. Alternatively, should a *S. aureus* strain carry a mutation within *ccpA* or its regulatory cascade, it is possible that glucose-dependent repression of TSST-1 could be lost. Therefore, this work highlights the necessity of understanding the dynamics of the vaginal environment as they relate to microbial composition, pH, and glucose.

When examining activation of PBMCs by *S. aureus* supernatants, we found that *S. aureus* supernatants grown in acidic conditions resulted in decreased IL-2 production (**Fig. 8**), despite an increase in TSST-1 production. While TSST-1 is the dominant superantigen produced by *S. aureus* MN8, the strain also encodes 7 additional superantigen genes which also contribute to the IL-2 profile (i.e. SEA and the enterotoxin gene cluster, *egc*). Their superantigenic activity is evidenced by the Δ*saeS* pH 6.8 supernatant still triggering IL-2 production from PBMCs, despite the lack of TSST-1 production in these conditions (**Fig. 8B**). Therefore, it is possible that the other *S. aureus* MN8 superantigens also have lower expression at pH 4.5, resulting in an overall decrease in T cell activation.

This acidic virulence surge that occurs at pH 4.5 begs the question of what its biological function may be. The ability of *S. aureus* to survive in an acidic vagina is curious, as this can be considered as a defense mechanism aimed at limiting vaginal pathogen survival. The eubiotic vagina ranges from pH 3.0–4.5, while *S. aureus* demonstrates deficits in growth *in vitro* below the range of 4.0–4.5 (**Fig. 2A-D**). While TSST-1 has often been associated with pathogenesis, our laboratory recently demonstrated in a murine vaginal colonization model that TSST-1 promotes colonization of the vaginal tract [28]. Another factor which promotes vaginal colonization are the *S. aureus* fibronectin-binding proteins [39], yet they have been found to be down-regulated in response to acidity, which may suggest a decrease in colonization potential [40]. Taken together, we propose that the biological function of TSST-1 overexpression at pH 4.5 may not necessarily amount to pathogenesis, rather it may be used as a colonization tactic in a harsh environment, particularly if other colonization factors are limited. Further studies examining pH, colonization versus pathogenesis, and TSST-1 production are required to elucidate this further.

The use of pAmilux in our vaginal colonization model serves as an important tool for measuring virulence gene expression *in vivo*. The use of luminescence reporters could be expanded to examine other genes of interests, as well as other sites of *S. aureus* colonization and infection, and for testing anti-virulent therapies *in vivo*. While we were able to identify GraXRS and SaeRS as activators of *tst* expression *in vivo* (**Fig. 9**), a limitation in our study is the pH of the mouse vaginal tract. The mouse vagina is not naturally colonized by lactobacilli, which in humans functions to acidify the vagina. Moreover, ‘humanized’ mice which are enriched in *Lactobacillus* dominated microbiotas, are also unable to attain a highly acidic vaginal pH (43). Consequently, we were unable to investigate the acidic virulence surge *in vivo* due to natural limitations. Further work is required within the field to develop animal models which better mimic the human female reproductive tract. Nevertheless, we have developed a method of monitoring bacterial gene expression during infection/colonization in real time.

Overall, this work highlights the importance of investigating a wide range of environmental conditions present in *S. aureus* niches when assessing pathogenesis. Our work highlights the potentially dangerous possibility of TSST-1 production occurring outside of menstrual conditions, with implications for menstrual health and safety. Altogether, our results demonstrate a complex sequence of events that occur in response to changes in pH and further indicate that environmental alterations may drastically alter the virulence potential of *S. aureus*.

## MATERIALS AND METHODS

### Ethics statement

For experiments with human PBMCS, healthy adult blood donors were recruited at the University of Western Ontario (UWO) and written, informed consent from each volunteer. Blood was anonymized, and no information regarding the identity of the donor, including sex and age, was retained as per the study protocol. The study was reviewed and approved by the UWO Health Sciences Research Ethics Board (HSREB #110859).

### Bacterial growth conditions

A list of bacterial strains used in this study can be found in **Table 1**. Wildtype and deletion mutants of *S. aureus* MN8 were plated on tryptic soy supplemented with 1.5 % agar (TSA). Strains containing plasmids were plated with the appropriate antibiotic concentrations. All *S. aureus* strains were grown overnight in tryptic soy broth (TSB), with or without antibiotics, at 37°C and 250 rpm unless otherwise specified. All *E. coli* strains were grown overnight in Luria-Bertani broth (LB), with or without antibiotics, at 37°C and 250 rpm unless otherwise specified. Vaginally defined medium (VDM) was prepared as previously described [10,23], and pH adjustments were performed with 1M HCl or 1M NaOH. Unless otherwise stated, VDM was formulated at 0.7 mM glucose.

**Table 1.** Bacterial strains used in this study.

| Strain | Description | Source |
| --- | --- | --- |
| *E. coli* XL1 blue | Cloning strain | Stratagene |
| *E. coli* SA30B | CC30 methylation strain | [41] |
| *S. aureus* MN8 | Prototypic mTSS strain | [42] |
| *S. aureus* MN8 Δ*graS* | Clean deletion of the *graS* sensor kinase in the two-component system GraXRS | This study |
| *S. aureus* MN8 Δ*saeS*Δ*graS* | Clean deletion of the *saeS* and *graS* sensor kinases in the two-component systems SaeRS and GraXRS respectively | This study |
| *S. aureus* MN8 Δ*ccpA* | Clean deletion of *ccpA* | [9] |
| *S. aureus* MN8 Δ*saeS* | Clean deletion of the *saeS* sensor kinase in the two-component system SaeRS | [6] |
| *S. aureus* MN8 Δ*sarA* | Clean deletion of *sarA* | [6] |
| *S. aureus* MN8 Δ*srrAB* | Clean deletion of the *srrAB* two-component system | [43] |
| *S. aureus* MN8 Δ*agr* | MN8 with the *agr* operon replaced by a *tetR* marker | [44] |
| *S. aureus* MN8 Δ*rot* | Clean deletion of *rot* | [4] |
| *S. aureus* MN8 Δ*agr*Δ*rot* | Clean deletion of *rot* and *agr* | [4] |
| *S. aureus* MN8 /pAmilux::*ptst* | MN8 containing the luminescent plasmid pAmilux under the control of the *tst* promoter; Cm10 | [44] |
| *S. aureus* MN8/pAmilux::*psaeP1* | MN8 containing the luminescent plasmid pAmilux under the control of the *sae P1* promoter; Cm10 | This study |
| *S. aureus* MN8/pAmilux::*psarA* | MN8 containing the luminescent plasmid pAmilux under the control of the *sarA* promoter; Cm10 | This study |
| *S. aureus* MN8/pAmilux::*pagrP3* | MN8 containing the luminescent plasmid pAmilux under the control of the *agr P3* promoter; Cm10 | This study |
| *S. aureus* MN8 /pAmilux::*pgraXRS* | MN8 containing the luminescent plasmid pAmilux under the control of the *graXRS* promoter; Cm10 | This study |
| *S. aureus* MN8 Δ*graS* /pAmilux::*ptst* | MN8 Δ*graS* containing the luminescent plasmid pAmilux under the control of the *tst* promoter; Cm10 | This study |
| *S. aureus* MN8 Δ*graS* /pAmilux::*psaeP1* | MN8 Δ*graS* containing the luminescent plasmid pAmilux under the control of the *sae P1* promoter; Cm10 | This study |
| *S. aureus* MN8 Δ*graS* /pAmilux::*pagrP3* | MN8 Δ*graS* containing the luminescent plasmid pAmilux under the control of the *agr P3* promoter; Cm10 | This study |
| *S. aureus* MN8 Δ*graS* /pAmilux::*psarA* | MN8 Δ*graS* containing the luminescent plasmid pAmilux under the control of the *sarA* promoter; Cm10 | This study |
| *S. aureus* MN8 Δ*saeS* /pAmilux::*ptst* | MN8 Δ*saeS* containing the luminescent plasmid pAmilux under the control of the *tst* promoter; Cm10 | [6] |
| *S. aureus* MN8 Δ*sarA* /pAmilux::*ptst* | MN8 Δ*sarA* containing the luminescent plasmid pAmilux under the control of the *tst* promoter; Cm10 | [6] |
| *S. aureus* MN8 Δ*srrAB*<br>/pAmilux::*ptst* | MN8 Δ*srrAB* containing the luminescent plasmid pAmilux under the control of the *tst* promoter; Cm10 | [45] |
| *S. aureus* MN8 Δ*ccpA*<br>/pAmilux::*ptst* | MN8 Δ*ccpA* containing the luminescent plasmid pAmilux under the control of the *tst* promoter; Cm10 | [9] |
| *S. aureus* MN8 Δ*agr*<br>/pAmilux::*ptst* | MN8 Δ*agr* containing the luminescent plasmid pAmilux under the control of the *tst* promoter; Cm10 | [44] |
| *S. aureus* MN8 Δ*rot*<br>/pAmilux::*ptst* | MN8 Δ*rot* containing the luminescent plasmid pAmilux under the control of the *tst* promoter; Cm10 | [4] |
| *S. aureus* MN8 Δ*agr*Δ*rot*<br>/pAmilux::*ptst* | MN8 Δ*agr*Δ*rot* containing the luminescent plasmid pAmilux under the control of the *tst* promoter; Cm10 | [4] |

### Western blotting

Western blots were performed as previously described [10]. Briefly, 6% trichloroacetic acid (TCA) precipitations were performed to concentrate exoproteins to 12 OD600 units and stored at -20°C. Samples were diluted with Laemmli buffer and run on 12% SDS-PAGE for 30 minutes at 85 V followed by 1 hour at 150 V. Gels were either stained with ReadyBlue, (Sigma-Aldrich) or transferred to a polyvinylidene difluoride membrane and blocked overnight in blocking buffer [10]. Western blotting was performed with 1:1,000 TSST-1 rabbit polyclonal anti-sera, and 1:20,000 IR-Dye800 goat anti-rabbit IgG antibody (Rockland). Blots were imaged using an ODESSY imager (LI-COR Biosciences).

### Construction of pAmilux plasmids

A list of primers used in this study can be found in **Table 2**. *S. aureus* promoters of interest were PCC amplified from *S. aureus* MN8 gDNA followed by a restriction digest and ligation with pAmilux. The ligated plasmid was then heat-shock transformed into *E. coli* XL1 blue (+Amp100), extracted, and then transformed into *E. coli* SA30B for appropriate methylation (+Amp100). One positive clone for each construct was extracted, sequenced, and then electroporated into competent *S. aureus* cells.

**Table 2.**
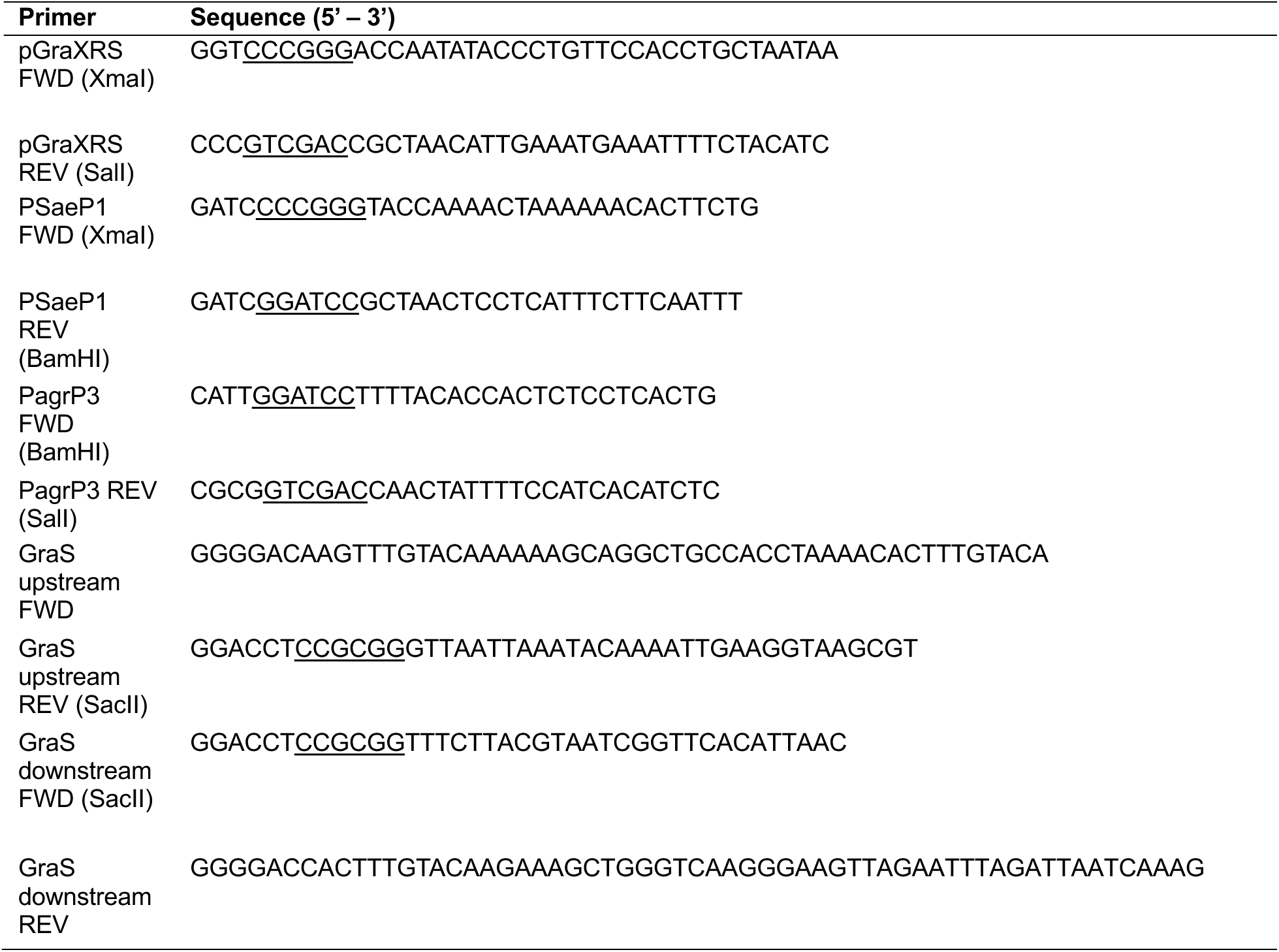
Primers used in this study.

### Luciferase Assays

Luciferase assays were performed as previously described [10]. Briefly *S. aureus* containing pAmilux reporter plasmids were grown overnight in TSB (+Cm10), followed by a three-hour subculture in VDM (pH 6.8, + Cm10). The cultures were then adjusted to an OD600 of 0.01 in the media of interest and plated in triplicate in a 96-well plate. The plate was incubated in a BioTek Synergy H4 reader at 37°C with continuous shaking on the medium setting. Data were plotted as the area under the curve (AUC) of the relative luminescence units, over the AUC of the OD600 readings. Data were analyzed using GraphPad Prism 10.

### Deletion of graS

The pKOR1 plasmid was used to create a markerless deletion of the *graS* gene as previously described [9,46]. Briefly, primers were created for the ∼800 bp regions upstream and downstream of *graS* and PCR amplified from wild-type MN8 gDNA (**Table 2**). The upstream and downstream PCR products were then ligated and recombined with pKOR1 (BP Clonase II, Thermo Fisher). The recombined plasmid was transformed into *E. coli* XL1 blue at 30°C (+Amp100), extracted, and then transformed into *E. coli* SA30B at 30°C for methylation (+Amp100). The methylated plasmid was extracted, sequenced, and electroporated into competent wild-type *S. aureus* MN8 or *S. aureus* MN8 Δ*saeS* cells at 30°C (+Cm10). The strains were grown in TSB at 30°C (+Cm10) overnight, then subcultured and grown overnight at 42°C (+Cm10). *S. aureus* clones were screened for integration and subcultured three times at 30°C prior to plating on ATc plates (1µg/mL). Colonies were picked and screened for chloramphenicol resistance, where sensitive colonies were then screened for the deletion, and sequenced.

### Peripheral blood mononuclear cell (PBMC) activation assays

PBMC assays were performed as previously described [10]. Briefly, *S. aureus* strains were grown overnight in VDM at pH 6.8 or pH 4.5. Supernatants were collected and stored at -20°C until use. Blood was collected from healthy blood donors and PBMCs were isolated via density gradient centrifugation. Cells (1 × 10^6^) were seeded and exposed to *S. aureus* supernatants. Cells were incubated overnight in complete RPMI [10], and the following day cell supernatants were collected and stored at -20°C. ELISAs were performed to detect IL-2 production (Invitrogen).

### Mice

Female BALB/cJ mice 7 to 9 weeks of age were used for *in vivo* experiments. Mice were purchased from Jackson Laboratory (stock 000651). Animals were housed without exceeding four animals per cage and provided water and food *ad libitum* with appropriate environmental enrichment. All animal experiments were in accordance with the Canadian Council on Animal Care Guide to the Care and Use of Experimental Animals, and the animal protocol was approved by the Animal Use Subcommittee at the University of Western Ontario (Protocol #2024-082).

### IVIS Vaginal Colonization Model

The vaginal colonization model was adapted from [28]. Briefly, 7- to 9-week-old female mice (Jackson Labs) (n=6) were provided with chloramphenicol water (2.25 mg/mL) *ad libitum* 24 hours prior to infection and refreshed every 24 hours, and pre-infection imaging was also performed (IVIS). Mice were vaginally inoculated with 1×10^6^ CFU of wild-type, Δ*graS,* or Δ*saeS* harboring pAmilux::p*tst*. Following inoculation, mice were imaged at 6, 8, 10, 12, and 24 hpi. At 24 hpi, the mice were sacrificed, and the lower reproductive tracts were excised, homogenized, and plated in the presence or absence of chloramphenicol for enumeration.

### Statistical Analysis

All statistical analysis was performed using GraphPad Prism 10. Ordinary one-way analysis of variance (ANOVA) was used without correction for multiple comparisons for luciferase data, IL-2 ELISAs and *in vivo* work. Unpaired t tests were also performed for luciferase data where specified.

## ACKNOWLEDGEMENTS

This project was funded by the Canadian Institutes of Health Research Project grant (PJT 198030) to J.K.M. C.S.M. was the recipient of an Ontario Graduate Scholarship (OGS) and a Guiding interdisciplinary Research On Women’s and girls’ health and Wellbeing (GROWW) Scholarship. N. R. W. was the recipient of an R.G.E. Murray graduate scholarship.

## SUPPORTING INFORMATION

**Figure S1.**
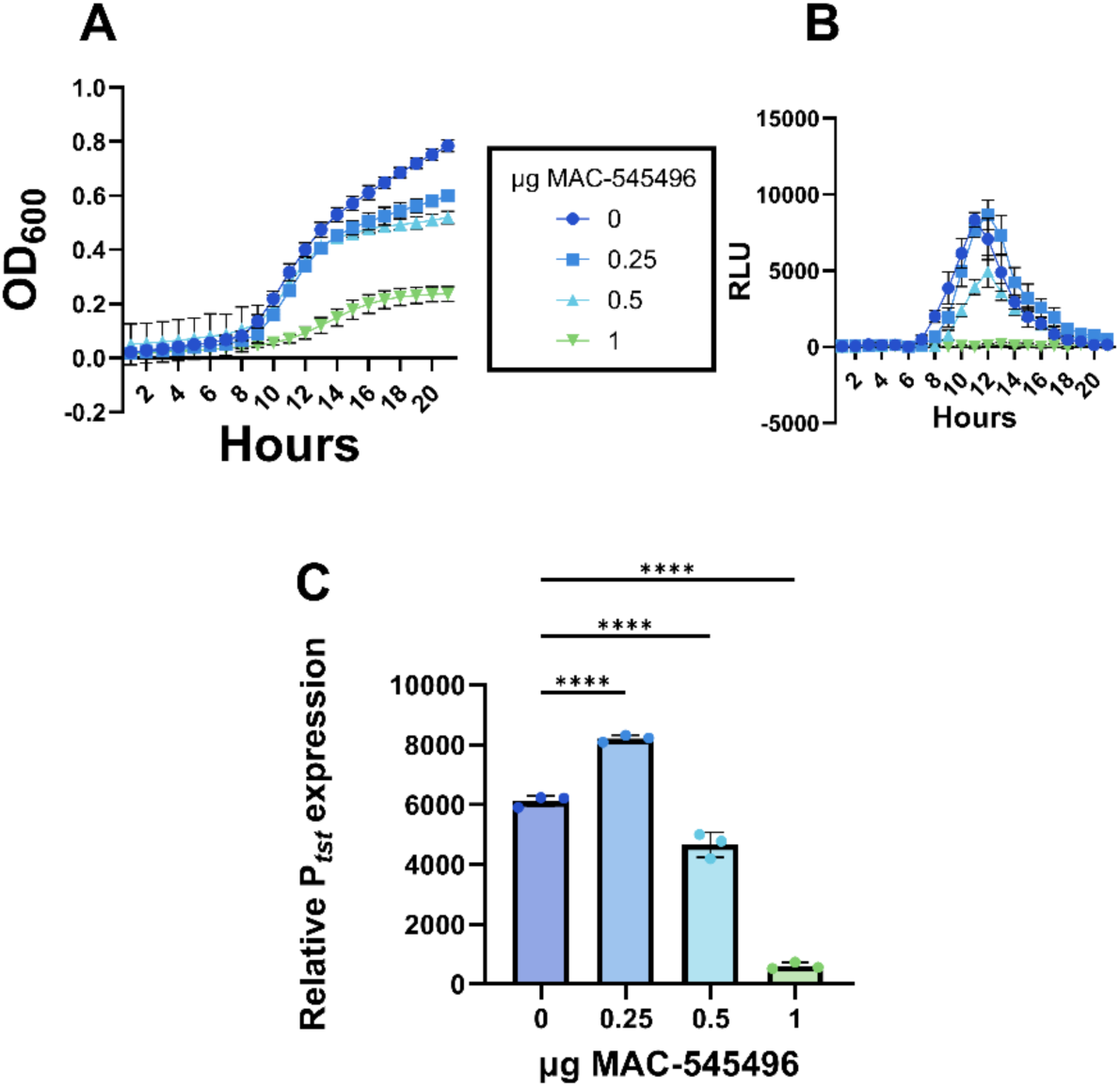
The GraR inhibitor MAC-545496 inhibits both growth and *tst* expression in *S. aureus* MN8. Wild-type MN8 containing pAmilux::p*tst* was grown in VDM at pH 4.5 with increasing amounts of the inhibitor MAC-545496 and readings were taken every hour (A, B). Relative expression was determined as the area under the curve (AUC) of luminescence over the AUC of growth. The uninhibited data is used as a comparison reference and is the same as in Fig. 2K. Statistical analysis was performed with GraphPad Prism 10 using a One-Way ANOVA (* P≤0.05, ** P≤0.01, *** P≤0.001, **** P≤0.0001).

**Figure S2.**
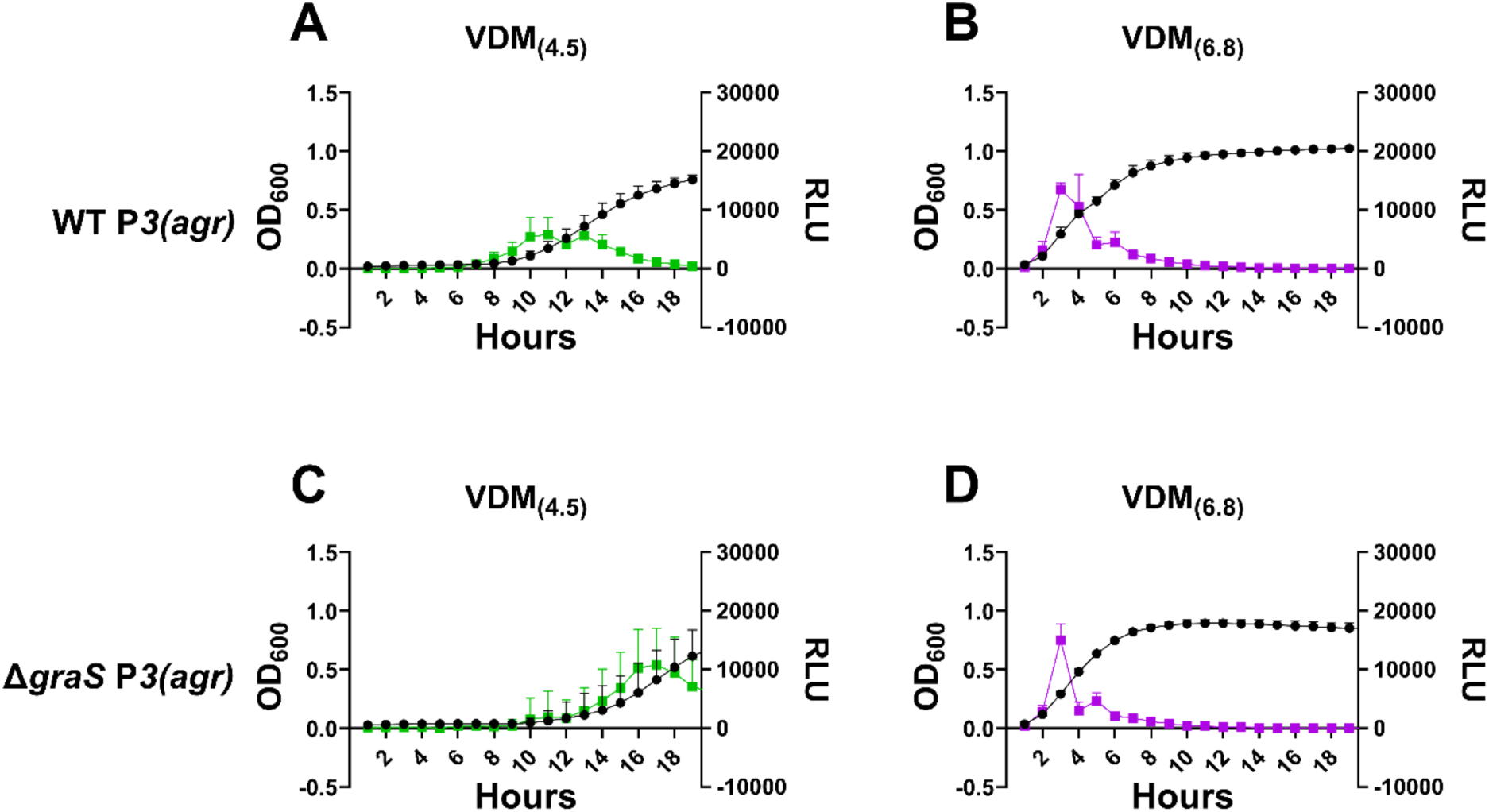
Expression from the *agr* P3 promoter is similar between wild-type MN8 and Δ*graS*. Wild-type MN8 (A, B) or MN8 Δ*graS* (C, D) containing pAmilux::p3*(agr)* were grown in VDM at pHs 4.5 and 6.8 and readings were taken every hour.

## Notes

### Competing Interest Statement

The authors have declared no competing interest.

## REFERENCES

1. Maduta CS, Tuffs SW, McCormick JK, Dufresne K. Interplay between *Staphylococcus aureus* and the vaginal microbiota. Trends in Microbiology. 2024;32: 228–230. doi:10.1016/j.tim.2023.12.005

2. Tuffs SW, Dufresne K, Rishi A, Walton NR, McCormick JK. Novel insights into the immune response to bacterial T cell superantigens. Nat Rev Immunol. 2024; 1–18. doi:10.1038/s41577-023-00979-2

3. Ali L, Abdel Aziz MH. Crosstalk involving two-component systems in *Staphylococcus aureus* signaling networks. Journal of Bacteriology. 2024;206: e00418–23. doi:10.1128/jb.00418-23

4. Tuffs SW, Herfst CA, Baroja ML, Podskalniy VA, DeJong EN, Coleman CEM, et al. Regulation of toxic shock syndrome toxin-1 by the accessory gene regulator in *Staphylococcus aureus* is mediated by the repressor of toxins. Mol Microbiol. 2019;112: 1163–1177. doi:10.1111/mmi.14353

5. Yarwood JM, McCormick JK, Schlievert PM. Identification of a Novel Two-Component Regulatory System That Acts in Global Regulation of Virulence Factors of *Staphylococcus aureus*. Journal of Bacteriology. 2001;183: 1113–1123. doi:10.1128/JB.183.4.1113-1123.2001

6. Baroja ML, Herfst CA, Kasper KJ, Xu SX, Gillett DA, Li J, et al. The SaeRS Two-Component System Is a Direct and Dominant Transcriptional Activator of Toxic Shock Syndrome Toxin 1 in *Staphylococcus aureus*. J Bacteriol. 2016;198: 2732–2742. doi:10.1128/JB.00425-16

7. Dufresne K, DiMaggio DA, Maduta CS, Brinsmade SR, McCormick JK. Discovery of an antivirulence compound that targets the *Staphylococcus aureus* SaeRS two-component system to inhibit toxic shock syndrome toxin-1 production. J Biol Chem. 2024;300: 107455. doi:10.1016/j.jbc.2024.107455

8. Sun F, Li C, Jeong D, Sohn C, He C, Bae T. In the Staphylococcus aureus two-component system sae, the response regulator SaeR binds to a direct repeat sequence and DNA binding requires phosphorylation by the sensor kinase SaeS. J Bacteriol. 2010;192: 2111–2127. doi:10.1128/JB.01524-09

9. Dufresne K, Podskalniy VA, Herfst CA, Lovell GFM, Lee IS, DeJong EN, et al. Glucose Mediates Niche-Specific Repression of *Staphylococcus aureus* Toxic Shock Syndrome Toxin-1 through the Activity of CcpA in the Vaginal Environment. Journal of Bacteriology. 2022;204: e00269–22. doi:10.1128/jb.00269-22

10. Maduta CS, McCormick JK, Dufresne K. Vaginal community state types (CSTs) alter environmental cues and production of the *Staphylococcus aureus* toxic shock syndrome toxin-1 (TSST-1). Journal of Bacteriology. 2024;206: e00447–23. doi:10.1128/jb.00447-23

11. O’Hanlon DE, Come RA, Moench TR. Vaginal pH measured in vivo: lactobacilli determine pH and lactic acid concentration. BMC Microbiol. 2019;19: 13. doi:10.1186/s12866-019-1388-8

12. Mirmonsef P, Hotton AL, Gilbert D, Burgad D, Landay A, Weber KM, et al. Free Glycogen in Vaginal Fluids Is Associated with *Lactobacillus* Colonization and Low Vaginal pH. PLoS One. 2014;9: e102467. doi:10.1371/journal.pone.0102467

13. Mirmonsef P, Hotton AL, Gilbert D, Gioia CJ, Maric D, Hope TJ, et al. Glycogen Levels in Undiluted Genital Fluid and Their Relationship to Vaginal pH, Estrogen, and Progesterone. PLoS One. 2016;11: e0153553. doi:10.1371/journal.pone.0153553

14. Amsel R, Totten PA, Spiegel CA, Chen KC, Eschenbach D, Holmes KK. Nonspecific vaginitis. Diagnostic criteria and microbial and epidemiologic associations. Am J Med. 1983;74: 14–22. doi:10.1016/0002-9343(83)91112-9

15. O’Hanlon DE, Moench TR, Cone RA. Vaginal pH and microbicidal lactic acid when lactobacilli dominate the microbiota. PLoS One. 2013;8: e80074. doi:10.1371/journal.pone.0080074

16. McCormick JK, Yarwood JM, Schlievert PM. Toxic shock syndrome and bacterial superantigens: an update. Annu Rev Microbiol. 2001;55: 77–104. doi:10.1146/annurev.micro.55.1.77

17. Weinrick B, Dunman PM, McAleese F, Murphy E, Projan SJ, Fang Y, et al. Effect of mild acid on gene expression in Staphylococcus aureus. J Bacteriol. 2004;186: 8407–8423. doi:10.1128/JB.186.24.8407-8423.2004

18. Bergdoll MS. Regulation and Control of Toxic Shock Syndrome Toxin 1: Overview. Reviews of Infectious Diseases. 1989;11: S142–S144.

19. Schlievert PM, Blomster DA. Production of staphylococcal pyrogenic exotoxin type C: influence of physical and chemical factors. J Infect Dis. 1983;147: 236–242. doi:10.1093/infdis/147.2.236

20. Flannagan RS, Kuiack RC, McGavin MJ, Heinrichs DE. *Staphylococcus aureus* Uses the GraXRS Regulatory System To Sense and Adapt to the Acidified Phagolysosome in Macrophages. mBio. 2018;9: e01143–18. doi:10.1128/mBio.01143-18

21. Kuiack RC, Veldhuizen RAW, McGavin MJ. Novel Functions and Signaling Specificity for the GraS Sensor Kinase of *Staphylococcus aureus* in Response to Acidic pH. J Bacteriol. 2020;202: e00219–20. doi:10.1128/JB.00219-20

22. Villanueva M, García B, Valle J, Rapún B, Ruiz de los Mozos I, Solano C, et al. Sensory deprivation in *Staphylococcus aureus*. Nat Commun. 2018;9: 523. doi:10.1038/s41467-018-02949-y

23. Geshnizgani AM, Onderdonk AB. Defined medium simulating genital tract secretions for growth of vaginal microflora. J Clin Microbiol. 1992;30: 1323–1326. doi:10.1128/jcm.30.5.1323-1326.1992

24. Anukam KC, Reid G. Effects of metronidazole on growth of *Gardnerella vaginalis* ATCC 14018, probiotic *Lactobacillus rhamnosus* GR-1 and vaginal isolate *Lactobacillus plantarum* KCA. Microbial Ecology in Health and Disease. 2008;20: 48–52. doi:10.1080/08910600701837964

25. Beetham CM, Schuster CF, Kviatkovski I, Santiago M, Walker S, Gründling A. Histidine transport is essential for the growth of *Staphylococcus aureus* at low pH. PLOS Pathogens. 2024;20: e1011927. doi:10.1371/journal.ppat.1011927

26. El-Halfawy OM, Czarny TL, Flannagan RS, Day J, Bozelli JC, Kuiack RC, et al. Discovery of an antivirulence compound that reverses β-lactam resistance in MRSA. Nat Chem Biol. 2020;16: 143–149. doi:10.1038/s41589-019-0401-8

27. Wilde AD, Snyder DJ, Putnam NE, Valentino MD, Hammer ND, Lonergan ZR, et al. Bacterial Hypoxic Responses Revealed as Critical Determinants of the Host-Pathogen Outcome by TnSeq Analysis of *Staphylococcus aureus* Invasive Infection. PLoS Pathog. 2015;11: e1005341. doi:10.1371/journal.ppat.1005341

28. Dufresne K, Al KF, Craig HC, Coleman CEM, Kasper KJ, Burton JP, et al. TSST-1 promotes colonization of *Staphylococcus aureus* within the vaginal tract by activation of CD8+ T cells. Infection and Immunity. 2025;0: e00439–24. doi:10.1128/iai.00439-24

29. Liu Q, Yeo W-S, Bae T. The SaeRS Two-Component System of *Staphylococcus aureus*. Genes (Basel). 2016;7: E81. doi:10.3390/genes7100081

30. Goerke C, Fluckiger U, Steinhuber A, Zimmerli W, Wolz C. Impact of the regulatory loci agr, sarA and sae of *Staphylococcus aureus* on the induction of alpha-toxin during device-related infection resolved by direct quantitative transcript analysis. Mol Microbiol. 2001;40: 1439–1447. doi:10.1046/j.1365-2958.2001.02494.x

31. Gao P, Wei Y, Hou S, Lai P-M, Liu H, Tai SSC, et al. SaeR as a novel target for antivirulence therapy against *Staphylococcus aureus*. Emerg Microbes Infect. 12: 2254415. doi:10.1080/22221751.2023.2254415

32. Falord M, Mäder U, Hiron A, Débarbouillé M, Msadek T. Investigation of the *Staphylococcus aureus* GraSR regulon reveals novel links to virulence, stress response and cell wall signal transduction pathways. PLoS One. 2011;6: e21323. doi:10.1371/journal.pone.0021323

33. Lindahl SH. Reviewing the options for local estrogen treatment of vaginal atrophy. International Journal of Women’s Health. 2014;6: 307. doi:10.2147/IJWH.S52555

34. Jenkins DJ, Woolston BM, Hood-Pishchany MI, Pelayo P, Konopaski AN, Quinn Peters M, et al. Bacterial amylases enable glycogen degradation by the vaginal microbiome. Nat Microbiol. 2023. doi:10.1038/s41564-023-01447-2

35. Nasioudis D, Beghini J, Bongiovanni AM, Giraldo PC, Linhares IM, Witkin SS. α-Amylase in Vaginal Fluid: Association With Conditions Favorable to Dominance of *Lactobacillus*. Reprod Sci. 2015;22: 1393–1398. doi:10.1177/1933719115581000

36. Nunn KL, Clair GC, Adkins JN, Engbrecht K, Fillmore T, Forney LJ. Amylases in the Human Vagina. mSphere. 2020;5: e00943–20. doi:10.1128/mSphere.00943-20

37. Spear GT, French AL, Gilbert D, Zariffard MR, Mirmonsef P, Sullivan TH, et al. Human α-amylase present in lower-genital-tract mucosal fluid processes glycogen to support vaginal colonization by *Lactobacillus*. J Infect Dis. 2014;210: 1019–1028. doi:10.1093/infdis/jiu231

38. Nunn KL, Ridenhour BJ, Chester EM, Vitzthum VJ, Fortenberry JD, Forney LJ. Vaginal Glycogen, Not Estradiol, Is Associated With Vaginal Bacterial Community Composition in Black Adolescent Women. J Adolesc Health. 2019;65: 130–138. doi:10.1016/j.jadohealth.2019.01.010

39. Lyon LM, Doran KS, Horswill AR. *Staphylococcus aureus* Fibronectin-Binding Proteins Contribute to Colonization of the Female Reproductive Tract. Infect Immun. 2023;91: e0046022. doi:10.1128/iai.00460-22

40. Costa FG, Mills KB, Crosby HA, Horswill AR. The *Staphylococcus aureus* regulatory program in a human skin-like environment. mBio. 2024;15: e00453–24. doi:10.1128/mbio.00453-24

41. Monk IR, Tree JJ, Howden BP, Stinear TP, Foster TJ. Complete Bypass of Restriction Systems for Major *Staphylococcus aureus* Lineages. mBio. 2015;6: 1–12. doi:10.1128/mBio.00308-15

42. Blomster-Hautamaa DA, Kreiswirth BN, Kornblum JS, Novick RP, Schlievert PM. The nucleotide and partial amino acid sequence of toxic shock syndrome toxin-1. J Biol Chem. 1986;261: 15783–15786.

43. Schlievert PM, Merriman JA, Salgado-Pabón W, Mueller EA, Spaulding AR, Vu BG, et al. Menaquinone analogs inhibit growth of bacterial pathogens. Antimicrob Agents Chemother. 2013;57: 5432–5437. doi:10.1128/AAC.01279-13

44. Li J, Wang W, Xu SX, Magarvey NA, McCormick JK. *Lactobacillus reuteri*-produced cyclic dipeptides quench *agr*-mediated expression of toxic shock syndrome toxin-1 in staphylococci. Proc Natl Acad Sci U S A. 2011;108: 3360–3365. doi:10.1073/pnas.1017431108

45. Tiwari N, López-Redondo M, Miguel-Romero L, Kulhankova K, Cahill MP, Tran PM, et al. The SrrAB two-component system regulates *Staphylococcus aureus* pathogenicity through redox sensitive cysteines. Proc Natl Acad Sci U S A. 2020;117: 10989–10999. doi:10.1073/pnas.1921307117

46. Bae T, Schneewind O. Allelic replacement in *Staphylococcus aureus* with inducible counter-selection. Plasmid. 2006;55: 58–63. doi:10.1016/j.plasmid.2005.05.005

